# Pre-trained molecular representations enable antimicrobial discovery

**DOI:** 10.1101/2024.03.11.584456

**Authors:** Roberto Olayo-Alarcon, Martin K. Amstalden, Annamaria Zannoni, Medina Bajramovic, Cynthia M. Sharma, Ana Rita Brochado, Mina Rezaei, Christian L. Müller

## Abstract

The rise in antimicrobial resistance poses a worldwide threat, reducing the efficacy of common antibiotics. Determining the antimicrobial activity of new chemical compounds through experimental methods is still a time-consuming and costly endeavor. Compound-centric deep learning models hold the promise to speed up this search and prioritization process. Here, we introduce a lightweight computational strategy for antimicrobial discovery that builds on `MolE`(`Mol`ecular representation through redundancy reduced `E`mbedding), a deep learning framework that leverages unlabeled chemical structures to learn task-independent molecular representations. By combining `MolE` representation learning with experimentally validated compound-bacteria activity data, we design a general predictive model that enables assessing compounds with respect to their antimicrobial potential. The model correctly identified recent growth-inhibitory compounds that are structurally distinct from current antibiotics and discovered *de novo* three human-targeted drugs as *Staphylococcus aureus* growth inhibitors which we experimentally confirmed. Our framework offers a viable cost-effective strategy to accelerate antibiotics discovery.

## Introduction

The development of novel antibiotics is a priority given the widespread dissemination of pathogenic strains resistant to current treatments [1]. Novel therapeutic candidates are often first identified by screening large chemical libraries. However, the success of these screenings is limited, with a typical hit rate between 1-3% [2, 3]. This issue is compounded by the high cost of the experiments owing to the large size of the chemical libraries being evaluated (from thousands to millions of molecules). Furthermore, the limited variability in these libraries makes it challenging to validate newly discovered or synthesized chemical species [2, 4]. In this context, deep learning-assisted strategies hold the promise to greatly contribute to the prioritization of molecules that should be experimentally evaluated, thus accelerating the rate at which novel drug candidates are found [5].

Using computational methods to estimate and predict properties of molecules, traditionally referred to as quantitative structure-activity relationships modeling, has a long history in material sciences, molecular biology, and biochemistry [6, 7]. The success of these modeling approaches hinges on an appropriate representation of the molecules, i.e., chemical descriptors such as the Extended Connectivity Fingerprint (ECFP) [8], and the availability of sufficiently large training data to map structure and activity. In recent years, the field of computational molecular property prediction has seen major breakthroughs, largely owing to advances in graph neural networks (GNNs) [9, 10, 11, 12]. These graph-based methods adopt an end-to-end supervised learning framework where predictive models infer a task-specific latent representation from large-scale training data. Important application benchmarks include the Tox21 data [13], where twelve toxic effects of 12,000 environmental chemicals and drugs were measured and provided as prediction challenges, and the MoleculeNet collection [14] which comprises prediction tasks for 16 datasets spanning different application domains. In the context of antimicrobial discovery, a major focus has been the prediction of antimicrobial peptides (AMPs) using curated databases such as CAMP_R3_ (see [15] and references therein). Despite the success of sophisticated deep-learning architectures for AMP prediction [16, 17, 18, 19], their task specificity limits the transferability of the learned representation to novel tasks and molecule types. Likewise, the application of Directed Message Passing Neural Networks (D-MPNNs) [12] for general antimicrobial discovery [2, 20, 21, 22] required the creation of a custom training set via in-house compound screening. A large number of compounds (ranging from 2,000 to 39,000 molecules) were tested for growth-inhibitory activity against each microbial species of interest, requiring considerable lab expertise and resources. Despite recent advances in lab automation and analysis [23], a publicly available large-scale data resource for generic antimicrobial discovery is not yet available, thus hindering the straightforward use of end-to-end learning schemes.

In this contribution, we tackle the challenge of antimicrobial discovery by introducing a two-stage deep-learning strategy that enables the assessment of the antibacterial potential of any compound of interest (Figure 1). The first stage uses a novel self-supervised pre-training strategy, termed `MolE` (`Mol`ecular representation through redundancy reduced `E`mbeddings), for molecule representation (Figure 1a). In the second stage we learn a set of antimicrobial potential (AP) scores for molecule prioritization that leverages the `MolE` representation and publicly available measurements of growth-inhibiting effects of FDA-approved drugs, including human-targeted and anti-infective drugs, against a diverse set of bacterial species [24] (Figure 1b and c). Inspired by recent molecular self-supervised learning schemes [25, 26, 27, 28], `MolE` leverages the large collection of available *unlabeled* chemical structures from PubChem [29] to learn a general-purpose molecular representation that is transferable to downstream prediction tasks. `MolE` uses Graph Isomorphism Networks (GINs) for representation learning and introduces the non-contrastive Barlow-Twins pre-training framework [30] to the molecular domain. Combined with supervised learning schemes, `MolE` enables competitive predictive performance on a series of curated molecular property prediction tasks from MoleculeNet.

**Figure 1:**
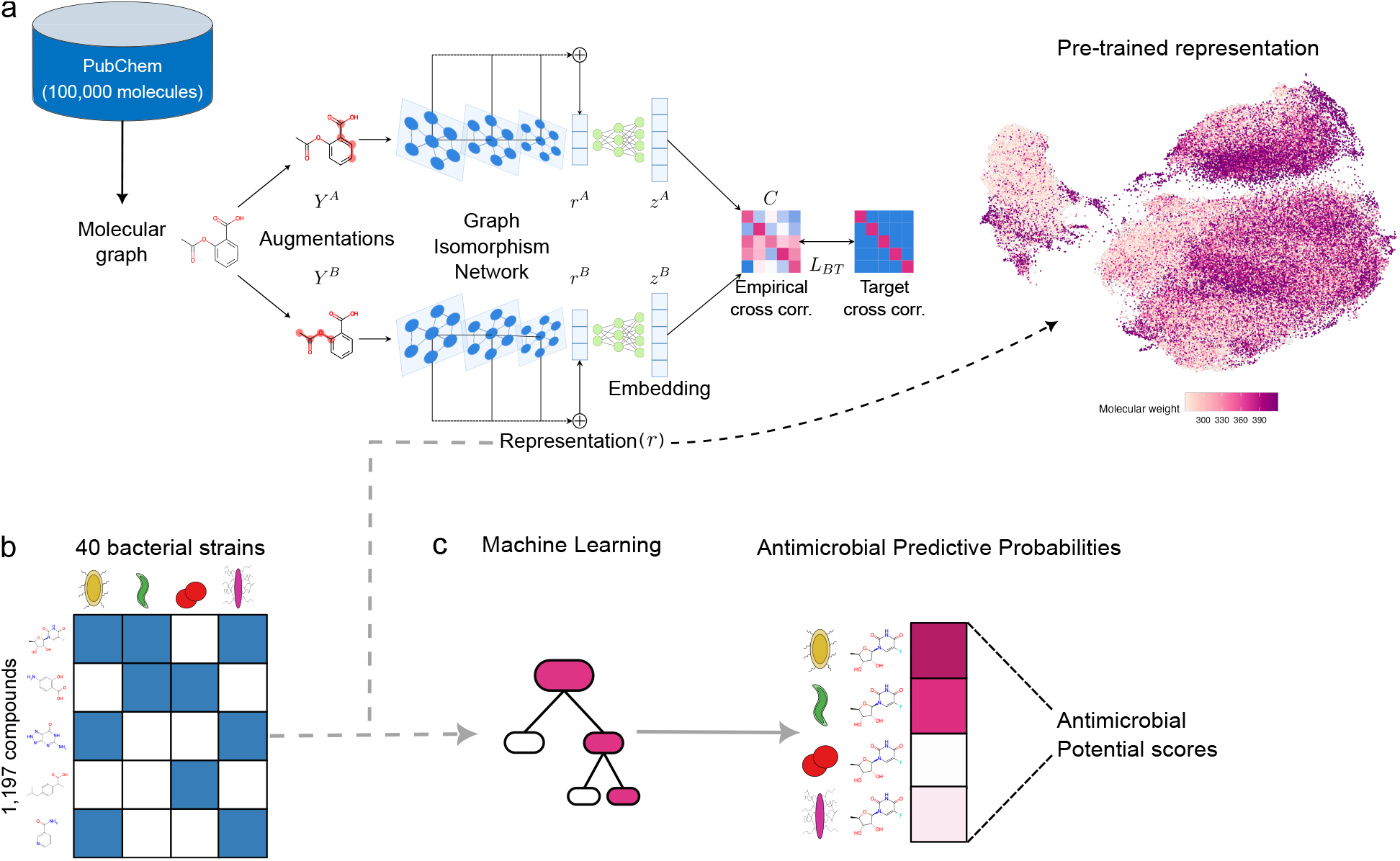
Two-stage framework for antimicrobial discovery. **a**. The `MolE` pre-training framework uses unlabeled structures from PubChem to learn a task-independent, molecular representation by optimizing the *L*_*BT*_ objective. **b**. Publicly available measurements of growth inhibition against 40 microbial strains [24] are used to train a predictive model. **c**. The pre-trained molecular representation is combined with the compound-microbe activity measurements to train a machine-learning model that produces a probability for each compound-microbe combination, indicating how likely the compound is to inhibit the microbe’s growth. These probabilities are used to estimate a collection of Antimicrobial Potential scores that serve to prioritize compounds for experimental validation.

For the purpose of antimicrobial discovery, we show that `MolE`-derived antimicrobial potential scores not only reflect the broad- and narrow-spectrum activity of structurally diverse compounds, such as Halicin [2] and Abaucin [31], but can also serve as compound prioritization guide in large-scale chemical screens. On a separate chemical library of over 2000 compounds, we identified approximately 200 compounds with high AP scores, many of which with yet to be discovered potential antibiotic activity. With our framework, we prioritized five compounds for experimental growth-assay validation on four bacterial species. We confirmed significant inhibitory effects of three of the five compounds on the growth of the human pathogen *Staphylococcus aureus*, reaching an excellent success rate compared to the state of the art [3].

We envision that the presented workflow and methodologies such as ours will be adopted by a wide range of microbiologists as a general and cost-effective way to prioritize and discover novel molecules with antibiotic properties.

## Results

### MolE learns meaningful compound representations

An important prerequisite for discovering compounds with potentially antimicrobial properties is the use of a general, yet efficiently tunable, representation of molecular structures. To provide such a context-independent representation we developed the `MolE` (`M`olecular representation through redundancy reduced `E`mbedding) framework (Figure 1a). `MolE` is a non-contrastive self-supervised deep learning scheme that constructs a representation of molecular structures by applying the Barlow-Twins redundancy reduction scheme [30]. The input into our pre-training framework is based on SMILES (simplified molecular-input line-input system [32]), a popular text-based representation of chemical structures. The workflow, illustrated in Figure 1a, comprises five main steps: (i) the SMILES representation of molecules is used to construct a molecular graph, where each node represents an individual atom and each edge a chemical bond in the molecule; (ii) two augmentations are created for each molecule by masking a subgraph of the original structure; (iii) batches of these augmentations enter a series of Graph Isomorphism Network (GIN) layers for feature extraction; (iv) the pooled output of each GIN layer is collected to form a final vector representation *r* ***∈*** R^*d*^; (v) a non-linear projection head expands each of the two vector representations into an embedding of higher dimensionality *z* ***∈*** R^*D*^, serving as input to the Barlow-Twins objective function *L*_*BT*_ (Methods). After pre-training, the static `MolE` representation *r* can readily serve as input for downstream machine learning applications (such as Figure 1b). Alternatively, concatenating the `MolE` architecture with an additional predictive layer allows fine-tuning of the pre-trained GIN layer parameters, thus making the representation adaptive to a specific downstream task.

To investigate `MolE`’s pre-trained representation, we first examined similarities between the representations of 100,000 test molecules not seen during pre-training. Figure 2a shows a UMAP [33] embedding of these molecules using the `MolE` representation. We observe that `MolE` learned similar representations for molecules with matching functional groups and/or related topological features. For instance, compounds consisting of a naphthalene group connected to a long carbon chain are placed closely in the embedding space (Figure 2a top middle). Other examples include Pyridines bound to central nitrogen heterocycles (Figure 2a top right), Benzenes surrounded by ether bonds (Figure 2a lower right), and Nitrogen heterocycles with various decorations (Figure 2a lower left). Notably, `MolE`’s representation also recognizes the similarities in the structure of short amino-acid chains (Figure 2a top left). Indeed, these peptides belong to a distinct cluster of molecules in the embedding space, indicating that `MolE` can distinguish this molecule type without any further fine-tuning. This offers an advantage over AMP-specific models by capturing broader chemical variability.

**Figure 2:**
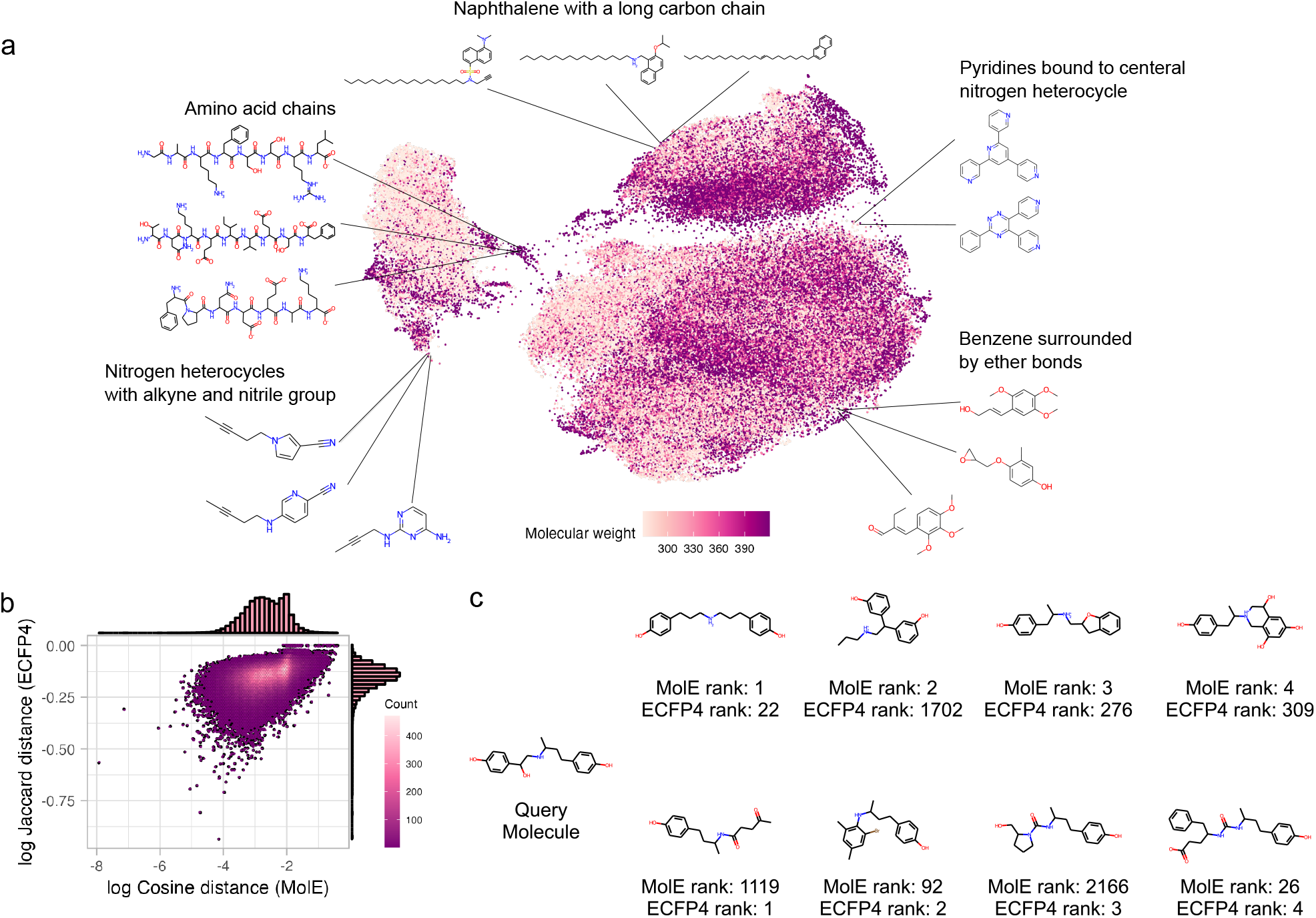
Illustration of `MolE`’s compound representation. **a**. UMAP embedding of `MolE`’s representation of 100,000 chemical structures not seen in pre-training. **b**. Comparison of Jaccard distance computed between ECFP4 representations and cosine distance computed between `MolE` representations with respect to the query molecule shown in **c**. Comparison of the four molecules with smallest distances (ranks) to the query according to `MolE` (top row) and ECFP4 (bottom row), respectively.

Compared to the standard ECFP4 representation, `MolE`’s representation captures distinct features for the same molecules. For illustration, we chose Ractopamine (PubChem ID: 56052, Figure 2c) as a hypothetical query molecule. We extracted its corresponding `MolE` and ECFP4 representations and calculated similarities to all other test set molecules (shown in Figure 2a) with the cosine and Jaccard distances, respectively (Methods). Figure 2b shows the high-level agreement between the two representations in terms of pair-wise distances to the query molecule (Spearman correlation: 0.42). However, discrepancies arise in the context of relevant nearest neighbors of the query, i.e., the most similar molecules to Ractopamine. Figure 2c ranks the top four most similar molecules in either representation. `MolE`’s most highly ranked molecules share two phenol groups connected by a carbon chain with one amine functional group with the query, ECFP4 only one phenol ring and a carbon chain with a methyl functional group. This illustrates `MolE`’s chemically meaningful embedding capabilities. Further examples are listed in Extended Data Figure 1, where `MolE` was able to highlight global structural features such as a naphthalene group connected to a long carbon chain (Extended Data Figure 1a), as well as the presence of specific functional groups such as sulfonyl chloride (Extended Data Figure 1b).

Taken together, these observations show that `MolE` captures chemically relevant information from unlabeled molecular structures. It recognizes the presence of functional groups and learns similar representations for molecules that share structural characteristics. The similarities recovered by `MolE` are distinct from those uncovered by ECFP4, potentially boosting the performance in downstream prediction tasks.

### MolE enables competitive molecular property prediction

To showcase `MolE`’s ability to achieve competitive molecular property prediction, we evaluated the out-of-sample performance of XGBoost [35] and Random Forests [36] when trained with `MolE`’s static representation as well as feed-forward neural networks that fine-tune `MolE`’s GIN layer weights. We considered selected classification and regression tasks from MoleculeNet [14] that are relevant for our context of compound prioritization in antimicrobial discovery, such as, e.g., the Tox21 and ClinTox benchmarks. Each dataset was split into training, validation, and testing sets following the scaffold splitting procedure [37], thereby creating a realistic scenario where molecules seen during training are structurally distinct from molecules seen during validation and testing. Performance is measured in terms of Area Under the Receiver Operating Characteristic curve (ROC-AUC) values obtained on the test set for each task.

We observed that `MolE`’s static representation (`MolE`_static_) combined with XGBoost outperformed alternative approaches in the majority of the classification and regression tasks (Table 1 and Extended Data Table 1, respectively). `MolE`_static_ enabled the best metric in four out of six classification tasks, with an average performance increase of 3% (in terms of average ROC-AUC) compared to the ECFP4 representation. Notably, XGBoost with `MolE` outperformed the end-to-end learning scheme D-MPNN in four classification tasks and two regression tasks. The added benefit of using the `MolE`_static_ representation was not limited to XGBoost as we observed similar performance gains with Random Forest predictors (Extended Data Tables 2 and 3). In general, fine-tuned `MolE`-based predictions (`MolE`_finetune_) displayed similar performance to their static counterparts. On the ClinTox classification benchmark, however, `MolE`_finetune_ achieved an average ROC-AUC of 92.85%, considerably outperforming all other approaches and even rivaling fine-tuned classifiers on large-scale ChemBERTa (90.6%) or MoLFormer (94.8%) representation models [25, 26]. This confirms that the computationally light-weight `MolE` framework can learn meaningful molecular representations with only *∼*100,000 unlabeled random structures from PubChem (Extended Data Figure 2d), starkly contrasting the large-scale pre-training schemes ChemBERTa [25] or MoLFormer [26], requiring millions of examples. Finally, the benchmarks also revealed `MolE`’s *non-contrastive* Barlow-Twins strategy to be superior to the state-of-the-art contrastive `MolCLR` framework. To identify the contributing factors of this performance improvement, we conducted an ablation study of all `MolE` components, including pretraining sample sizes and representation dimensions. (for details see Methods section and Extended Data Figure 2).

**Table 1:** Average ROC-AUC (%) and standard deviation obtained on classification benchmark tasks. The first 5 models are supervised learning methods. The next 4 are the names of molecular features given as input to an XGBoost classifier. The final four methods are fine-tuned models. The best performance metric for each category is marked in **bold**.

| Dataset | BBBP | Tox21 | ClinTox | BACE | SIDER | HIV |
| --- | --- | --- | --- | --- | --- | --- |
| # Molecules | 2039 | 7831 | 1478 | 1513 | 1427 | 41127 |
| # Tasks | 1 | 12 | 2 | 1 | 27 | 1 |
| GCN [27] | 71.8±0.9 | 70.9±2.6 | 62.5±2.8 | 71.6±2.0 | 53.6±3.2 | 74.0±3.0 |
| GIN [27] | 65.8±4.5 | 74.0±0.8 | 58.0±4.4 | 70.1±5.4 | 57.3±1.6 | **75.3±1.9** |
| SchNet [27] | 84.8±2.2 | **77.2±2.3** | 71.5±3.7 | 76.6±1.1 | 53.9±3.7 | 70.2±3.4 |
| MGCN [27] | **85.0±6.4** | 70.7±1.6 | 63.4±4.2 | 73.4±3.0 | 55.2±1.8 | 73.8±1.6 |
| D-MPNN [27] | 71.2±3.8 | 68.9±1.3 | **90.5±5.3** | **85.3±5.3** | **63.2±2.3** | 75.0±2.1 |
| ECFP4 | 68.51±1.01 | 70.76±1.48 | 84.52±0.00 | **84.33±1.12** | 62.58±1.89 | 75.77±1.6 |
| N-Gram | 74.00±0.01 | 73.51±0.54 | **89.96±1.27** | 81.97±0.01 | 63.68±1.59 | **78.89±0.48** |
| MolCLR<sub>static</sub> | 65.74±0.68 | 72.05±1.34 | 76.04±3.05 | 68.78±0.00 | 62.68±2.40 | 74.29±0.01 |
| MolE<sub>static</sub> | **75.98±0.32** | **76.15±0.61** | 84.90±0.01 | 83.84±0.86 | **64.48±2.65** | **78.52±0.01** |
| Hu et.al [27] | 70.8±1.5 | **78.7±0.4** | 78.9±2.4 | **85.9±0.7** | 62.7±0.8 | **80.20±0.9** |
| HiMol [34] | 73.2±0.80 | 76.2±0.30 | 80.8±1.4 | 84.6±0.20 | 62.5±0.30 | 74.71±0.23 |
| MolCLR<sub>finetune</sub> | 73.17±2.13 | 74.61±1.67 | 87.79±5.31 | 81.96±1.11 | 63.23±3.05 | 77.61±0.92 |
| MolE<sub>finetune</sub> | **75.34±0.93** | 75.21±1.77 | **92.85±1.33** | 83.21±0.93 | **64.98±4.42** | 78.98±0.57 |

Taken together, the benchmarking results indicate that (i) `MolE` produces a molecular representation that enables excellent downstream prediction performance even when pre-trained on a small dataset of 100,000 unlabeled structures (Extended Data Figure 2d) and (ii) `MolE` is particularly competitive for downstream tasks with a small number of labeled data (such as ClinTox and BACE).

### MolE enables generalizable predictions of antimicrobial compound activity against human gut microbes

The second stage of our framework (see Figure 1b and c) leveraged `MolE`’s representation capabilities to learn a set of antimicrobial potential scores that allow ranking and prioritization of compounds in the antimicrobial discovery process. To this end, we made use of the publicly available dataset created by Maier et al. [24], which evaluated the effect of 1,197 marketed drugs on the growth of 40 bacterial strains representative of the human gut microbiome (Figure 1b). This dataset was used to train an XGBoost model to learn the probability of growth inhibition for all available compound-microbe pairs (Figure 3a and b). For any compound of interest, the trained model, termed `MolE`-XGBoost, delivers a 40-dimensional vector of predictive probabilities (Figure 3b). Data-driven thresholding of these probabilities allows for (i) binary classification of a compound to be growth-inhibitory of a specific species or (ii) assessment of narrow- or broad-spectrum activity of the compound by inspecting the overall number of inhibited strains, respectively (Figure 3c, Equation 10).

**Figure 3:**
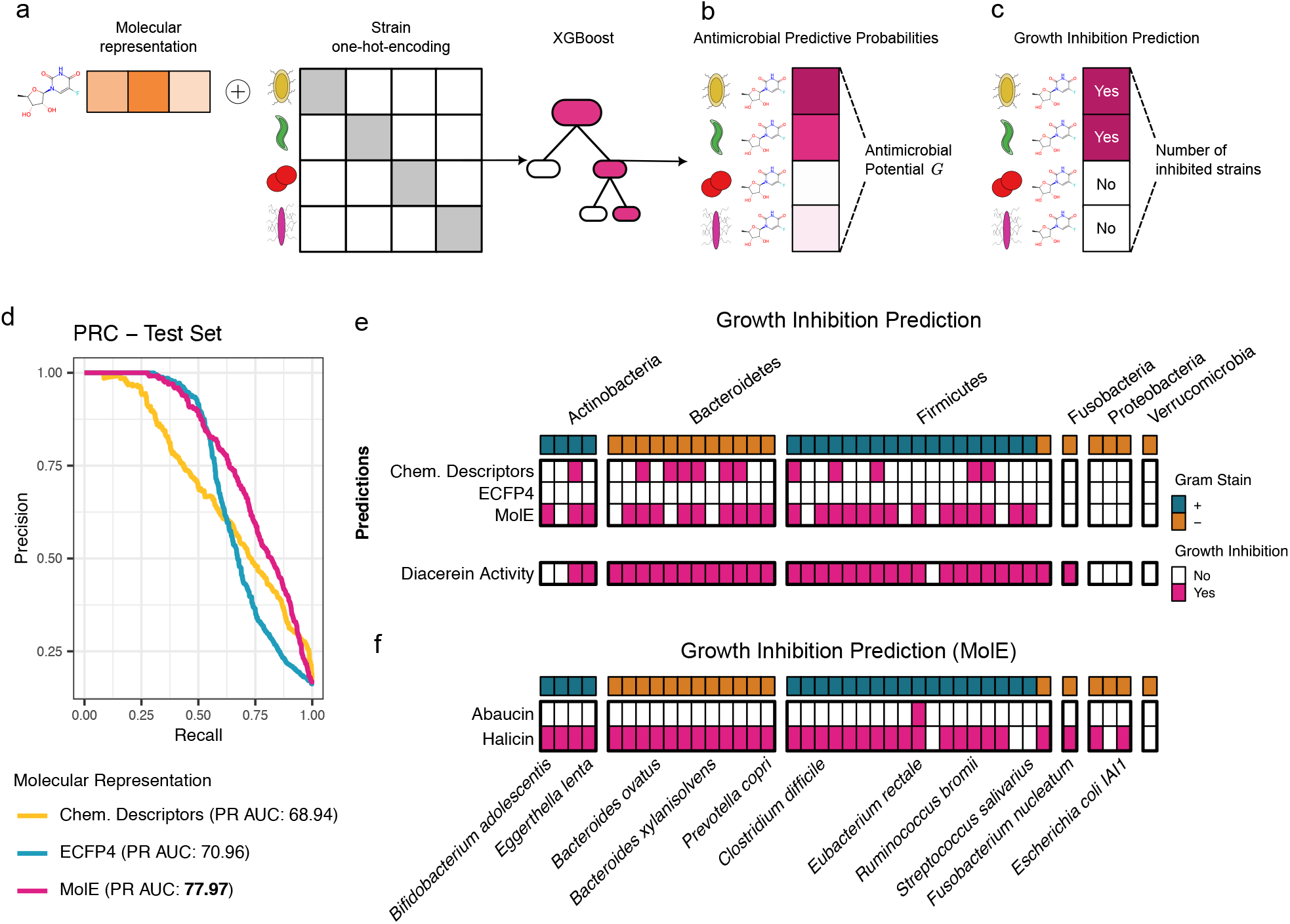
Predicting antimicrobial activity in the human gut microbiome. **a**. The molecular representation (ECFP4, Chemical Descriptors or `MolE`) is concatenated with a one-hot-encoding of the microbial strains present in the training dataset to train an XGBoost model. **b**. The model produces an antimicrobial predictive probability for each compound-microbe pair. The log-geometric mean of all probabilities corresponds to AP score *G*. **c**. The antimicrobial predictive probabilities are thresholded into binary predictions of growth inhibition; then, the total number of strains predicted to be inhibited is determined. **d**. Precision-recall curves obtained on the test set by models trained with ECFP4, Chemical Descriptors, and `MolE` representations. PR-AUC is shown in legend. **e**. Binary predictions for the growth-inhibitory activity of Diacerein made by models trained with Chemical Descriptors, ECFP4, or `MolE`. The experimentally validated activity of Diacerein is shown in the bottom row. Each column is an individual strain. **f**. Predictions for the antimicrobial activity of Halicin and Abaucin made by `MolE`-XGBoost.

`MolE`-XGBoost strongly outperformed ECFP4-based models and a recently proposed predictive model [4] that uses a collection of explicit chemical descriptors in terms of Precision and Recall (Figure 3d). Furthermore, `MolE`-XGBoost was the only model to predict the broad-spectrum activity of the human-targeted drug Diacerein, which was a part of the test set in our experiment (Figure 3e). The model correctly predicted an effect on 25 of the 33 strains that showed inhibited growth in lab experiments (Figure 3e bottom row).

Overall, our test set contained 24 compounds experimentally determined to have broad-spectrum antimicrobial activity, inhibiting the growth of 10 or more strains [24]. Of these, five compounds were uniquely recovered by `MolE`-XGBoost (Extended Data Figure 3). We did not observe examples of broad-spectrum compounds recovered by ECFP4- or Chemical Descriptor-based models that were not detected by `MolE`-XGBoost.

We further confirmed the generality of `MolE`-XGBoost by recapitulating the findings from recent ground-breaking studies that identified structurally novel antibiotic candidates [2, 31]. Firstly, our `MolE`-XGBoost model was able to re-discover the broad-spectrum activity of Halicin [2] (Figure 3f, bottom row) and correctly predicted the experimentally validated inhibition of *Escherichia coli* and *Clostridium difficile*. Secondly, in the case of Abaucin [31], the model recovered its highly narrow-spectrum activity, predicting only the (yet to be tested) inhibition of *Eubacterium rectale* (Figure 3f bottom row).

### Predicting antimicrobial potential in an orthogonal chemical library

Next, we used `MolE`-XGBoost’s predictions to identify *de novo* bacterial growth inhibitors in an orthogonal chemical library of 2,327 FDA-approved drugs, food homology products, and human endogenous metabolites.

We used the model’s predictive probabilities with respect to the 40 bacterial strains to design four variants of Antimicrobial Potential (AP) scores that allow the ranking of individual compounds: (i) the total number of strains predicted to be inhibited (Figure 3c), (ii) the log-geometric mean of estimated probabilities across all 40 strains (*G*, Figure 3b, Equation 11), (iii) the log-geometric mean of the probabilities of all 22 gram-positive strains (*G*^+^, Equation 12), and (iv) the log-geometric mean of the probabilities of all 18 gram-negative strains (*G*^*−*^, Equation 13).

Figure 4a illustrates the diversity of the library’s 2,327 compounds in a `MolE`-based UMAP embedding, highlighting compounds predicted to inhibit the growth of 10 or more strains. Among these, 158 compounds (*∼* 6.8% of the chemical library) were non-antibiotics. Our extensive post-hoc literature review revealed that, out of these 158 compounds, 53 had previously been reported to inhibit the growth of various bacterial species (Figure 4b). Examples include structurally diverse compounds such as natural products Ellagic acid [38] and Shionone [39], as well as human-targeted drugs such as Doxifluridine [40, 41] and Ospemifene [42] (Figure 4a). This putative 33% hit rate is an improvement over the 1-3% rate commonly cited for large-scale chemical screens [3].

**Figure 4:**
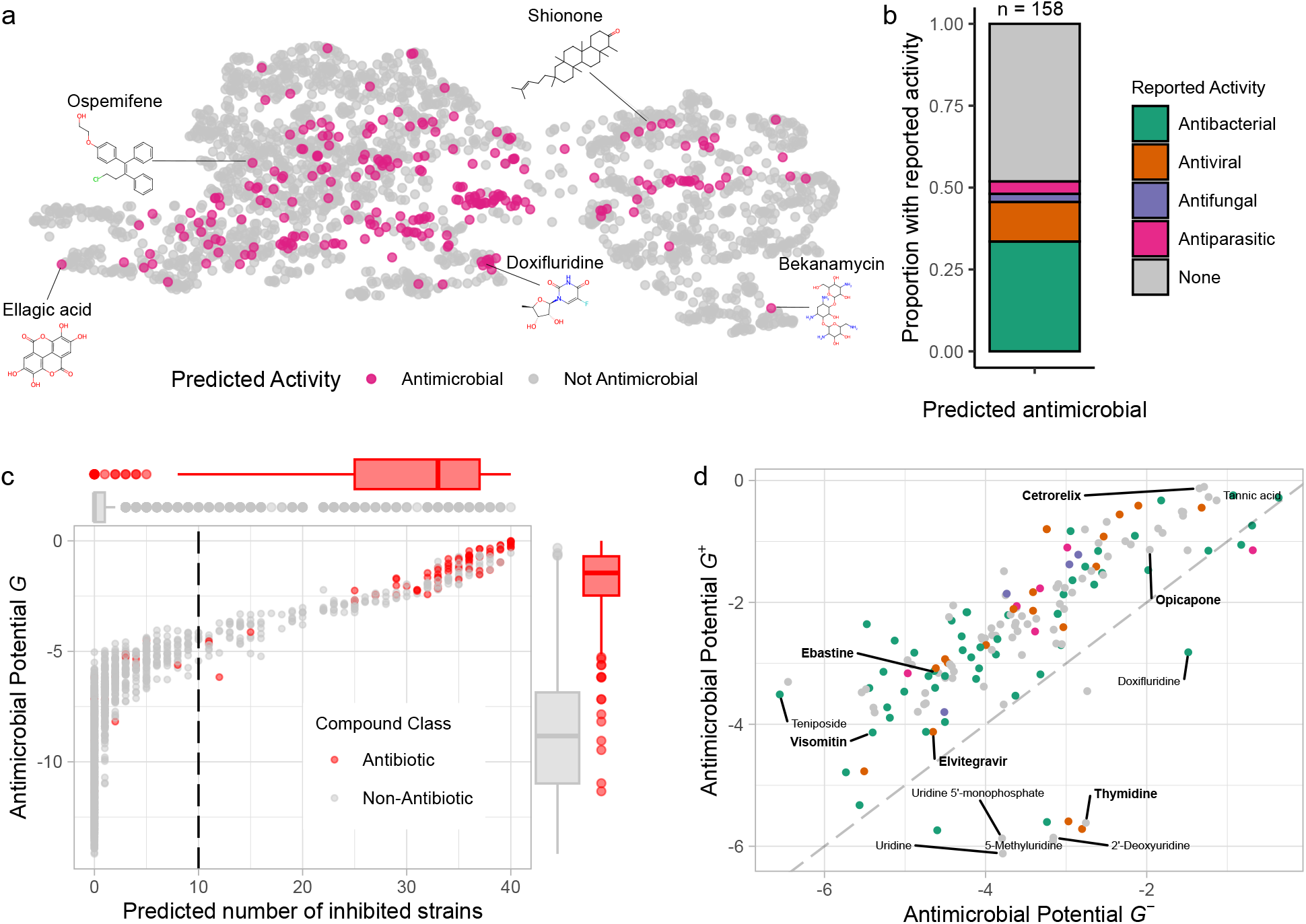
Predicting antimicrobial potential in a new chemical library. **a**. UMAP embedding of `MolE`’s representation of the 2,327 compounds for which predictions are made. Compounds predicted to inhibit 10 or more strains are highlighted in pink. **b**. Literature-reported activity of 158 non-antibiotic compounds predicted to inhibit at least 10 strains. **c**. Ranking of compounds based on Antimicrobial Potential scores. The predicted number of inhibited strains is compared to Antimicrobial Potential *G*. Known antibiotics are shown in red. **d**. Comparison of the Antimicrobial Potential for Gram-positive (*G*^+^) and Gram-negative strains (*G*^*−*^) determined for non-antibiotic drugs with predicted broad-spectrum activity. Names shown in bold were selected for experimental validation. Color legend is same as in panel **b**.

We further assessed the validity of our proposed AP scores by testing their ranking capabilities on the 93 antibiotics present in the chemical library. Figure 4c shows the distribution of general AP score *G* vs. the predicted number of inhibited strains for all compounds (Figure 3c). We observed remarkable ranking capability, with strong enrichment of antibiotic compounds at large AP scores (Figure 4c). Furthermore, 77 of the 93 antibiotics were predicted to inhibit 10 or more strains (Figure 4c, dashed line), confirming that both scoring schemes generalize well to unseen antibiotic compounds.

Finally, we assessed the discriminative potential of the refined AP scores for Gram-positive and Gram-negative strains by simultaneously ranking the 158 non-antibiotic compounds previously predicted to inhibit 10 or more strains (Figure 4d). In line with current knowledge about the susceptibility of Gram-positive bacteria to chemical stressors [24, 43, 44], we confirmed that most compounds exhibited greater AP scores against Gram-positive strains compared to Gram-negative strains (Figure 4d, above the dashed line). However, we observed that nucleotide analogs, such as Uridine and Uridine derivatives, were predicted to be more active against Gram-negative strains (Figure 4d, below the dashed line), confirming recent evidence that Uridine molecules are powerful adjuvants for the activity of aminoglycosides against *Escherichia coli* [45].

### Experimental validation of predicted antibacterial compounds

Guided by the AP scores on the orthogonal chemical library, we next selected five compounds to be experimentally validated via a minimum inhibitory concentration assay: Elvitegravir, Opicapone, Cetrorelix, Thymidine, and Visomitin. The selected compounds (i) were predicted to inhibit at least 10 strains, (ii) were commercially available, and (iii) covered a variety of functions and chemical structures (Figure 4d, shown in bold). Elvitegravir, Opicapone, Cetrorelix, and Thymidine had not been previously reported to inhibit the growth of bacterial or fungal strains. As a positive control, we included Visomitin as a non-antibiotic drug with proven antimicrobial activity [46]. We screened these compounds against a panel of bacterial strains that included the four human pathogens *Escherichia coli* UTI, *Klebsiella pneumoniae, Pseudomonas aeruginosa*, and *Staphylococcus aureus*, all of which were *not* present in the dataset by Maier et al. We also included one shared commensal strain, namely *E. coli* IAI1 (Figure 5a).

**Figure 5:**
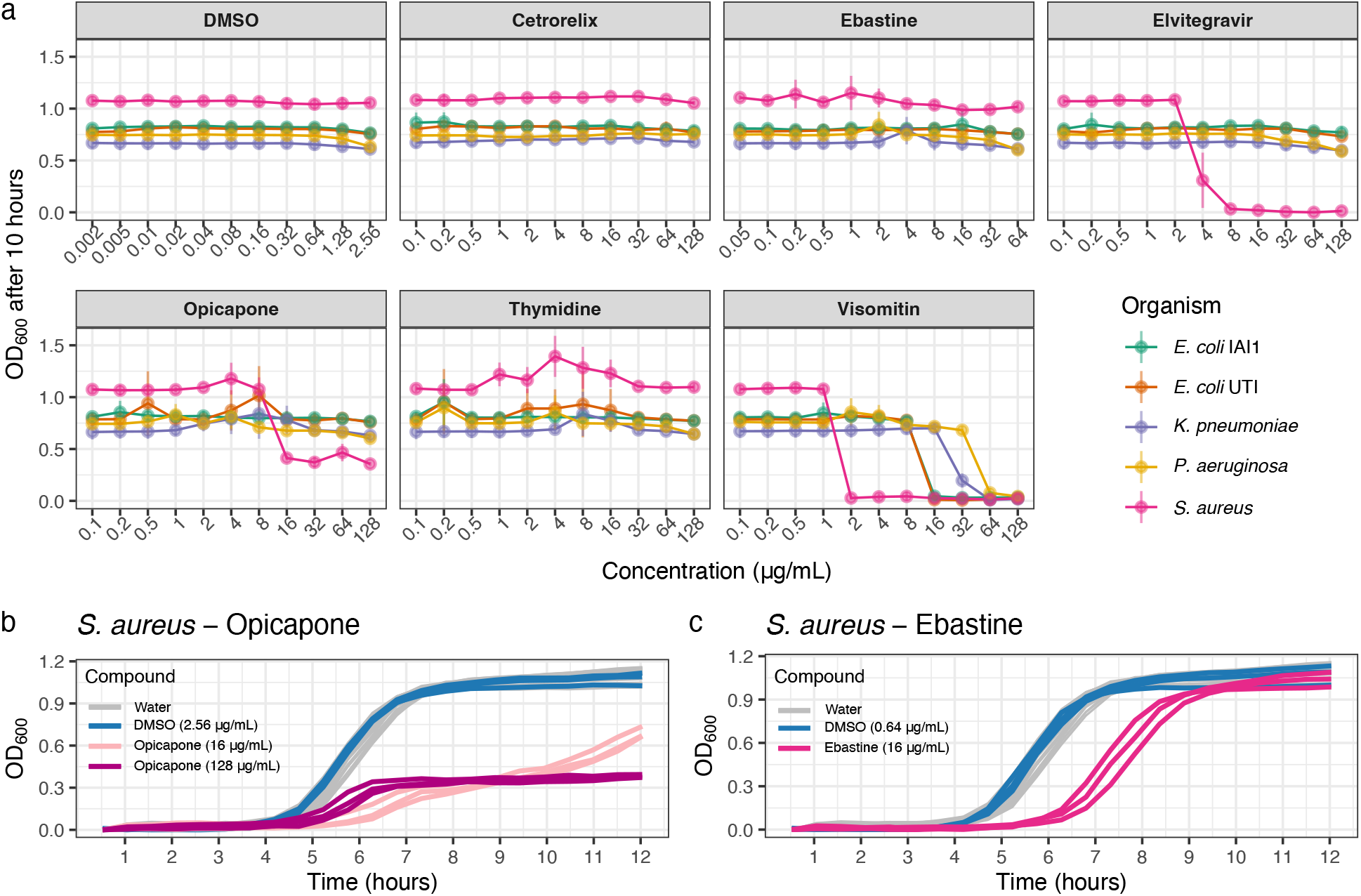
Experimental validation of antimicrobial activity. **a**. Average OD_600_ measurements (*±* standard deviation) after 10 hours of growth at increasing concentrations of each compound. **b**. Growth curves for *S. aureus* when grown in the presence of Water, DMSO, and Opicapone. **c**. Growth curve for *S. aureus* when grown in the presence of Water, DMSO, and Ebastine.

Overall, three of the five tested compounds were confirmed to have measurable effects on the growth of the Gram-positive pathogen *S. aureus*. The strongest effect was observed for Elvitegravir, which inhibited the growth of *S. aureus* at a concentration of 8 *µ*g/mL (Figure 5a). Furthermore, Opicapone significantly limited the growth of *S. aureus* to a maximum optical density of 0.38 *±* 0.01 at a concentration of 128 *µ*g/mL (Figure 5b). At a lower concentration of 16 *µ*g/mL, it extended both the lag phase duration and the population doubling time by about 1 hour compared to growth observed with DMSO exposure (Extended Data Figure 4b and c). Finally, Ebastine extended the duration of the lag phase of *S. aureus* by approximately 2 hours (Figure 5c and Extended Data Figure 5c). Notably, our experiments re-discovered the broad-spectrum activity of Visomitin, showing that this compound can inhibit the growth of *P. aeruginosa* and *K. pneumoniae* at 64 *µ*g/mL, thereby expanding the list of species known to be susceptible to this compound [46].

While we did not observe growth inhibition in response to Cetrorelix and Thymidine, their predicted Antimicrobial Potential can be explained by the presence of similar molecules that affect the growth of bacteria in the training set such as, e.g., Azidothymidine. This medication is a chemical analog of Thymidine and was previously shown to inhibit the growth of twelve strains (nine Gram-negative and three Gram-positive) [24]. As a result, Thymidine was predicted to be particularly effective against Gram-negative strains by our model (Figure 4d). For comparison, we also included the predicted antimicrobial effects of ECFP4- and Chemical Descriptor-based models for the five compounds in Extended Data Figure 6a-f.

In summary, the experimental validation demonstrated a remarkable discovery rate of our proposed *MolE* - based workflow, given the small number of compounds screened. The varied effects on bacterial growth observed for each compound suggest different mechanisms of action that can be further investigated in future studies.

## Discussion

In this contribution, we have presented (i) a computationally lightweight, predictive end-to-end workflow to identify novel antimicrobial candidates and (ii) experimentally confirmed growth-inhibitory effects of several compounds at a notable hit rate. Our framework specifically addresses the pervasive scarcity of biological and chemical data in the antimicrobial discovery process by leveraging the vast amounts of unlabeled chemical structures to learn a novel molecular representation within the `MolE` framework (Figure 1a). This pre-trained representation captures relevant chemical and structural features (Figure 2) and improves the performance of downstream machine learning algorithms when predicting molecular properties when few labeled examples are available (Table 1 and Extended Data Table 1). The ability to obtain competitive performance from `MolE`’s representation enables microbiology researchers with variable access to high-performance computing resources to make meaningful predictions of their property of interest, thus helping to democratize research molecular property prediction in microbiology and beyond.

We have shown that by using `MolE`’s pre-trained representation, standard XGBoost prediction models enable concise assessment of the antimicrobial potential of chemical compounds on publicly available data (Figure 3). Our `MolE`-XGBoost model re-discovered *de novo* structurally distinct antibiotic candidates such as Halicin and recognized the broad-spectrum activity of other compounds that would have been missed by standard models. By designing and validating a set of model-derived antimicrobial potential (AP) scores, we have been able to provide a reliable ranking of the growth-inhibiting potential of any compound of interest, not only for specific microbial strains but also for the class of Gram-positive or Gram-negative bacteria, and for bacteria in general (Figure 4 and Figure 5). This allows a much broader applicability of our framework in the antibiotics discovery process.

While large-scale (blind) experimental screening of chemical libraries will remain a common technique for drug discovery, it is unlikely to keep pace with the continuous introduction of novel chemical species. Therefore, methodologies such as ours are a practical alternative to prioritize molecules for screening. We have demonstrated these potential efficiency gains with the experimental validation of our model predictions, where three out of five compounds were found to have significant effects on the growth of *S. aureus* (Figure 5). These effects ranged from total growth inhibition to delays in the onset of exponential growth and slower growth rates. Screening additional strains, particularly more Gram-positive strains, could further uncover the growth-inhibitory activity of a greater number of predicted broad-spectrum antimicrobials [24]. While the compounds evaluated in this study require further research to be re-purposed into new antibiotic treatments, they are interesting study subjects when considering the effect of human-targeted drugs on microbial life. This is especially relevant, given the growing body of research showing that non-antibiotic drugs contribute to the appearance of bacterial strains resistant to current antibiotics [47, 48, 49, 50]. Future work can investigate the molecular mechanisms of action of these compounds, leading to a better understanding of the effect non-antibiotic drugs have on microbial growth.

We consider our proposed framework as a first important step toward the more general goal of computationally-guided antimicrobial discovery in the face of data scarcity. We envision that several future research directions can improve the present framework. For example, the interpretability of `MolE` representation may be enhanced by enabling quick identification of molecular characteristics associated with properties of particular interest for microbiology. Furthermore, alternative deep-learning architectures such as graph-attention networks or graph-transformer networks [51] may improve downstream prediction tasks. Incorporating biological features of microbial strains into the representation learning process may also help improve the specificity of predictions, enabling the identification of effective narrow-spectrum treatments [4].

In conclusion, our proposed framework addresses key challenges in the field of antimicrobial discovery. By overcoming the need for large data resources, we envision that the presented workflow and methodologies such as ours are better poised to complement large screenings, thus increasing the rate at which new treatment candidates are uncovered.

## Methods

### Pre-training

#### Dataset

The `MolE` pre-training dataset was created by randomly sampling 100,000 unlabeled molecules from a collection of 10,000,000 unique structures originally collected by ChemBERTa [25]. This subset is then randomly split into training (90%) and validation (10%).

#### Molecular graph construction

Each molecule is represented as a graph *G*, where all atoms are nodes *V* and the chemical bonds between them are the edges *E*. The attributes encoded for each atom are the element it represents and its chirality type (tetrahedral CW, tetrahedral CCW, other). Likewise, each bond is embedded by its type (single, double, triple, aromatic), and its direction (none, end-upright, end-downright).

As shown in Figure 1a, during pre-training two augmentations (*Y* ^*A*^ and *Y* ^*B*^) are created by following the subgraph removal procedure, first described in MolCLR [27]. Briefly, a seed atom is selected at random. This seed atom and its neighbors are masked and then neighbors of the neighbors are masked until 25% of the atoms present in the original have been masked. The bonds between these masked atoms are removed, producing a subgraph of the original molecule.

#### Graph Neural Networks

We explored the ability of Graph Isomorphism Networks (GINs) and Graph Convolutional Networks (GCNs) to extract meaningful features from the molecular graphs constructed in the previous step. Both algorithms use an aggregation function to learn an updated representation of each node. In the case of GINs, aggregation depends on an MLP head:

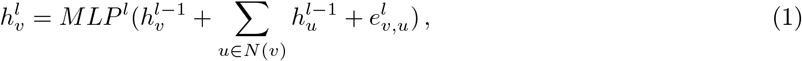

where 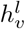 is the updated representation of node *v* at the *l*th GIN layer, *N* (*v*) indicates the neighbors of *v*, and *e*_*v,u*_ represents the embedding of the attributes of the bond between *v* and its neighbor *u*.

#### Molecular representation

To obtain a global vector representation of the molecular structure, we first gather a graph-level representation for each layer (*g*^*l*^).

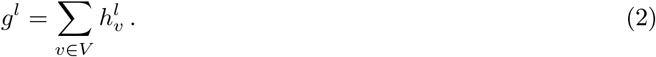

A final graph representation *r* is obtained by concatenating the graph-level representations of each layer into a single vector.

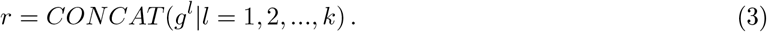

Here, *k* represents the total number of GNN layers used. In our setup, the dimensionality of *g*^*l*^ is set as a 200-dimensional vector. Given that *k* = 5, *r* is therefore a 1000-dimensional vector. In experiments where no concatenation is performed, *g*^*l*^ is set as a 1000-dimensional vector, and *r* = *g*^*k*^. The graph representation learned for augmentation *Y* ^*A*^ is correspondingly denoted as *r*^*A*^, as is the representation learned for augmentation *Y* ^*B*^ denoted as *r*^*B*^.

#### Non-contrastive learning

Once the final graph representations *r*^*A*^ and *r*^*B*^ are produced, an MLP layer is used to obtain embeddings *z*^*A*^ and *z*^*B*^, which are *d*-dimensional vectors. These embedding vectors are used to evaluate the *L*_*BT*_ objective function [30]. First, an empirical cross-correlation matrix *C* is computed between *z*^*A*^ and *z*^*B*^:

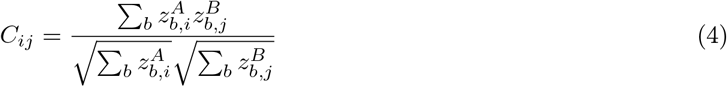

Here, *b* indexes the batch samples and *i* and *j* index the vector dimensions of the embeddings. In practical terms, *C* is a *d* x *d* matrix and represents the average cross-correlation matrix of a given batch *b*. Finally, *C* is used to calculate the *L*_*BT*_ loss:

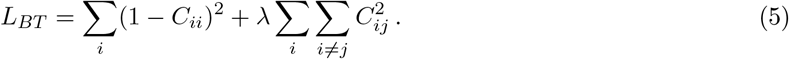

In essence, by minimizing *L*_*BT*_ we minimize the difference between *C* and the identity matrix. In this way, embeddings obtained from augmentations of the same molecule are made to be invariant to distortions, while at the same time penalizing redundancy of information encoded by the components of the embedding. The positive constant *λ* controls the trade-off between these two terms.

#### Pre-training setup

For all experiments, `MolE` pre-training is done over 1000 epochs with a batch size of 1000 molecules. An Adam optimizer with weight decay 10^*−*5^ is used to minimize *L*_*BT*_. The first 10 epochs are performed with a learning rate of 5 x 10^*−*4^, after which cosine learning decay is performed. The parameter configuration that minimizes the validation error is kept for downstream tasks.

### Benchmarking

#### Datasets

We collected 11 benchmark datasets originally curated and made available by MoleculeNet [14]. This set included 6 binary classification tasks and 5 regression tasks. To make these benchmarking scenarios more challenging and realistic, we split each dataset into training (80%), validation (10%), and test (10%) sets following the scaffold splitting procedure described by Hu et.al [37].

#### Training and evaluation

In our benchmarking experiments, `MolE` was pre-trained on 100,000 molecules from Pubchem [29]. Different dimensionalities of the embedding vectors (*z*^*A*^ and *z*^*B*^) and values for the trade-off parameter *λ* were explored, while the dimensionality of the representation *r* is fixed as a 1000-dimensional vector. After pre-training, we evaluated the resulting representation in two ways: 1) the static vector representation *r* was used as input for machine-learning algorithms (termed `MolE`_static_), or 2) the weights learned by the GIN layers were used to fine-tune a predictor for a specific task (termed `MolE`_finetune_).

In the first case, the pre-trained representation `MolE`_static_ was used as input molecular features to Random Forest or XGBoost algorithms. Hyperparameter optimization is performed via random search (details in Supplementary Information). Each model configuration was trained three times with different seed values. The model configuration with the largest mean ROC-AUC value on the validation set was then evaluated on the test set.

In the case of fine-tuning, an untrained MLP head is placed after the GNN layers. For all classification tasks, the Softmax cross-entropy loss is optimized, while in the case of regression, the *L*_1_ loss is optimized for the QM7 and QM8 tasks and the mean squared error is optimized for all other tasks. A random search is performed for hyperparameter optimization (Supplementary Information). The selected architecture was trained for 100 epochs, using an Adam optimizer with a weight decay of 10^-6.^ While the GNN and the MLP are updated during training, the learning rate chosen for both parts differed. Each model configuration was trained three times.

Extended Connectivity Fingerprints (ECFP) were calculated using the `GetMorganFingerprintsAsBitVect` function available in RDKit 2020.09.1.0 [52]. In order to get ECFP4 fingerprints we set `fp_radius`=2 and `fp_bits`=1024.

#### Other predictors

In Table 1 and Extended Data Table 1, performance metrics for GCN, GIN, SchNet, MGCN, D-MPNN, and Hu et al. are taken from the publication of MolCLR [27]. The ROC-AUC values for HiMol were taken from the respective publication [34], except for the HIV task which is evaluated in the current study.

The MolCLR_GIN_ [27] model made available on their GitHub page was used for all benchmarks, following the instructions in the same repository. The molecular representation obtained after GNN feature extraction was used as the input for either XGBoost or Random Forest. The N-Gram [53] and HiMol [34] models were pre-trained and molecular features were extracted following the default instructions and parameters made available in their respective GitHub repositories.

### Ablation study on the MolE framework

To identify the components of `MolE` that contribute to performance gains, we performed an ablation study shown in Extended Data Figure 2.

#### GNN backbone and construction

We compared GINs and GCNs in our graph feature extraction step combined with two alternatives for constructing the molecular representation *r*. Concatenated representations are obtained as described previously (Equation 3). Non-concatenated representations consisted of the pooled output of the last GNN layer. We observed that models perform best when trained on representations built with the concatenated strategy, independent of the GNN backbone used during pre-training (Extended Data Figure 2a). This indicates that each GNN layer captures important information about the molecular structure. In the ClinTox task, both GIN-derived representations outperformed their GCN counterparts.

#### Embedding dimensionality, trade-off parameter, and dataset size

We observed increased performance on the ClinTox task when *z*^*A*^ and *z*^*B*^ were 8000-dimensional vectors (Extended Data Figure 2b). We also found performance increases as the *λ* trade-off parameter approaches 10^*−*4^ (Extended Data Figure 2c). Finally, we noted that the size of the unlabeled dataset used during pre-training does not necessarily improve performance on most classification tasks (Extended Data Figure 2d). A higher ROC-AUC value can be observed in the ClinTox task when `MolE`’s representation is pre-trained on 200,000 molecules.

With these observations, we decided to pursue the task of antimicrobial discovery using the `MolE`_static_ representation obtained after pre-training on 100,000 molecules with *λ* = 10^*−*4^, *z*^*A,B*^ ∈ ℝ^8000^, using a GIN backbone and constructed by concatenating graph-layer representations.

### Exploring the MolE representation

#### Representation similarity

The distance between two `MolE` representations *r*^*i*^ and *r*^*j*^ was determined using the cosine distance:

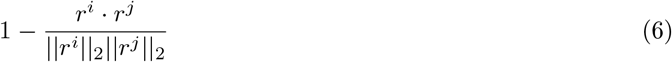

At the same time, ECFP4 vectors were compared by determining the Jaccard distance between two binary vectors.

#### Uniform Manifold Approximation and Projection

A UMAP embedding based on the cosine distance between `MolE` representations was built using the `umap-learn` 0.5.3 Python module.

### Predicting antimicrobial activity on human gut bacteria

#### Dataset

The adjusted p-value table from Maier et al [24] was used to determine labels for the growth-inhibitory effects of the screened compounds. Compound-microbe combinations with an adjusted p-value *<* 0.05 were considered to be examples of growth inhibition. The 1,197 molecules were divided into training (80%), validation (10%), and test (10%) sets following the scaffold splitting procedure.

#### Molecular and bacterial strain representation

We used the SMILES string of a given molecule *i* to obtain the corresponding representation *m*_*i*_, which is a *d*-dimensional vector. In our work, *m* can be one of three possible representations: i) ECFP4, in which case *d* = 1024, ii) `MolE`_static_ (*d* = 1000), and iii) a set of explicit Chemical Descriptor features (*d* = 92), described by Algavi and Borenstein [4], which were calculated using RDKit.

A given microbial strain *j* ∈ *B* (where *B* is the complete set of bacterial strains) is represented as a one-hot-encoded vector *b*_*j*_. Given that 40 strains are present in the dataset, *b*_*j*_ is a 40-dimensional vector.

#### Compound-microbe predictions of antimicrobial activity

The vectors *m*_*i*_ and *b*_*j*_ are concatenated to form *x*_*ij*_, which is a 40 + *d*-dimensional vector. This combination of molecular and microbe representations is the input to our classifier function *f* (*·*). In our work, *f* (*·*) is an XGBoost model that, for each *x*_*ij*_, outputs a probability (*p*_*ij*_) that indicates the likelihood of compound *i* inhibiting the growth of microbe *j*.

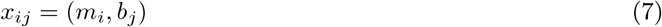

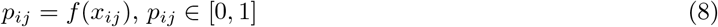

#### Ranking compounds by predicted antimicrobial activity

We summarise the predicted spectrum of activity of compound *i* two ways: i) As the total number of strains predicted to be inhibited by the compound (*K*_*i*_). This is achieved by thresholding the antimicrobial predictive probabilities (*p*_*ij*_) into binary predictions of growth inhibition and adding up the number of strains predicted to be inhibited.

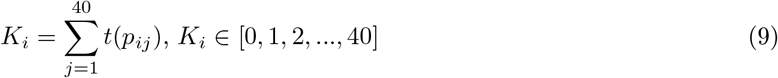

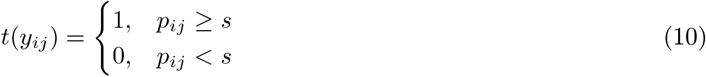

Here, the function *t*(*·*) binarizes the output by determining whether *p*_*i*_*j* exceeds a threshold *s*. We selected *s* to optimize the F1-Score metric obtained on the validation set for each model. This optimized cutoff was 0.044, 0.068, and 0.209 for the model trained with `MolE`, Chemical Descriptors, and ECFP4 features respectively.

ii) We introduce the Antimicrobial Potential score for compound *i* (*G*_*i*_), which is the log geometric mean of the antimicrobial predictive probabilities obtained for all microbes:

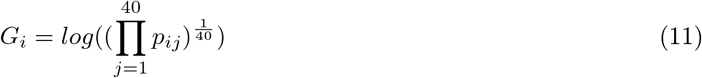

Additionally, we consider the value of the Antimicrobial Potential score when calculated on the subsets of gram-positive 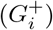 and gram-negative 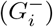 microbes. In Equations 12 and 13 we impose a fixed indexing on the set of taxa, where the first 22 indices represent all gram-positive bacteria (*j* ∈ [1, 2, …, 22]) and the remaining 18 indices represent all gram-negative bacteria (*j* ∈ [23, 24, …, 40])

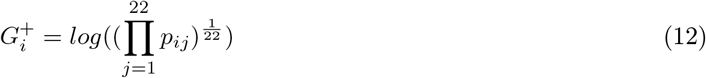

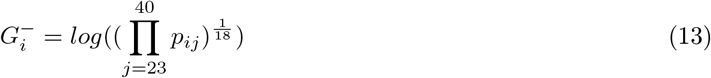

#### Model selection and evaluation

A random search over XGBoost hyperparameters was performed for each chemical representation. The model configuration with the highest PR-AUC on the validation set was then evaluated on the test set.

### Predicting antimicrobial compounds in an orthogonal chemical library

#### Chemical library

A separate chemical library was constructed based on the FDA-approved Drugs, Human Endogenous Metabolite, and Food-Homology Compound libraries made available by MedChemExpress https://www.medchemexpress.com/. The chemical structures for these compounds were gathered from PubChem [29] using the `pubchempy` 1.0.4 module. SMILES were canonicalized and salts were removed using the `SaltRemover` function from RDKit [52].

#### Compound annotation

Information provided by MedChemExpress included descriptions of the chemicals in the library. The corresponding Anatomical-Therapeutic-Chemical (ATC) code was assigned to each compound by matching compound name strings. A complete collection of ATC codes was taken from https://github.com/fabkury/atcd. Overlap with chemicals in the library used by Maier et al.[24] was determined by matching chemical names and ATC codes. Chemicals present in both libraries were not considered for downstream prediction.

#### Prediction and evaluation

Molecules with *K*_*i*_ *≥* 10 were prioritized for further evaluation. We performed a literature search for articles available on PubMed [54] that described the *in-vitro* and/or *in-vivo* antimicrobial activity of our prioritized compounds against any bacterial species. The results of our literature search and the complete set of predictions can be found in our GitHub repository (https://github.com/rolayoalarcon/mole_antimicrobial_potential).

### Experimental validation

#### Compound prioritization

In total 6 compounds were selected for experimental validation. Criteria considered for compound selection were the following: 1) The compound was predicted to inhibit 10 strains or more, 2) the compound was not an antibiotic, 3) the compound did not have antifungal activity, 4) no previous literature describing the antimicrobial activity of the compound was found, and 5) the compound could be purchased through an independent provider. Furthermore, we attempted to choose compounds with different biological functions that were structurally diverse from each other.

#### Bacterial strains and growth conditions

Before the experiments, all strains were cultured overnight in Lysogeny Broth (LB Lennox) adjusted to pH 7.5 at 37^*o*^C. Detailed information about the strains used in this study can be found in Supplementary Information.

#### Measurement of Minimum Inhibitory Concentrations (MIC)

All compounds were purchased from Biomol GmbH (Germany). Stock solutions were prepared in DMSO and stored at *−*20^*o*^C till further use. MIC measurements were performed in 96-well plates with 100 *µ*L bacterial cultures in Mueller Hinton (MH) broth using 1:2 serial dilutions of the tested compounds. Starting concentrations of 128 *µ*g/mL were used for Cetrorelix, Opicapone, Thymidine, Visomitin, and Elvitegravir, and 64 µg/mL for Ebastine due to low solubility. No-compound controls contained DMSO. Overnight cultures of three biological replicates of each bacterial strain were adjusted to OD_600_ = 0.1 and inoculated into the plates by pinning using a Singer Rotor (Singer Instruments, UK), achieving a 1:200 dilution. Plates were sealed with transparent breathable membranes (Breathe-Easy®, Sigma-Aldrich-Merck) and incubated at 37 °C in a Cytomat 2 incubator (Thermo Scientific) with continuous shaking at 800 rpm. OD_600_ was measured at regular 30-minute intervals for up to 12 h in a Synergy H1 plate reader (Agilent, USA).

#### Growth curve modelling

All growth curves were normalized by subtracting the minimum of the second, third, and fourth measurements taken. Afterward, each individual curve was modeled as a logistic curve with `Sicegar` 0.2.4 R package. From these curves, parameters such as the maximum growth rate *µ*_max_, end of lag-phase, start of stationary phase, and the carrying capacity were extracted. The maximum doubling time *t*_d_ was estimated as:

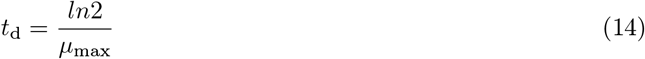

### Software and reproducibility

The `MolE` pre-training framework was implemented using the pytorch-geometric 1.6.3 framework [55] with Python 3.7 [56]. We use the `RandomForestClassifier` and `RandomForestRegressor` implementation available in scikit-learn 1.0.2 [57] and the `XGBClassifier` and `XGBRegressor` objects from xgboost 1.6.2 [58]. The scikit-learn 1.0.2 [57] module is also used when computing ROC-AUC, PR-AUC, and F1 score metrics. The R 4.3.1 language was used for its `ggplot2` 3.4.2 for plotting, `Sicegar` 0.2.4 packages. ECFP4, chemical descriptors, and general SMILES processing were done with the rdkit 2020.09.1.0 package.

## Data and Code Availability

The unlabeled chemical structures used for pre-training were gathered from the MolCLR GitHub repository https://github.com/yuyangw/MolCLR. The adjusted p-value table from Maier et al., 2018 [24] was used to train models to predict antimicrobial activity.

The code for the MolE pre-training framework can be found at https://github.com/rolayoalarcon/MolE. The steps and models for predicting antimicrobial activity are available at https://github.com/rolayoalarcon/mole_antimicrobial_potential.

## Acknowledgements

This work was funded by a grant awarded to C.L.M., A.R.B. and C.M.S. for the StressRegNet consortium within the Bavarian research network bayresq.net funded through the Bavarian State Ministry of Science and Arts, Germany.

## Author contributions

R.O.A. and C.L.M. conceived the overall objectives and design of the project. R.O.A., C.L.M. and M.R. contributed to the development and evaluation of the pre-training framework. R.O.A., A.R.B., C.M.S. A.Z. and M.K.A. conceived the experimental validation of antimicrobial activity for selected compounds. A.Z. and M.K.A. performed experiments. R.O.A. and M.B. analyzed data from experimental validations. R.O.A. implemented all computational methods. All authors revised and approved the final version of the Article.

## Extended Data

**Extended Data Table 1:**
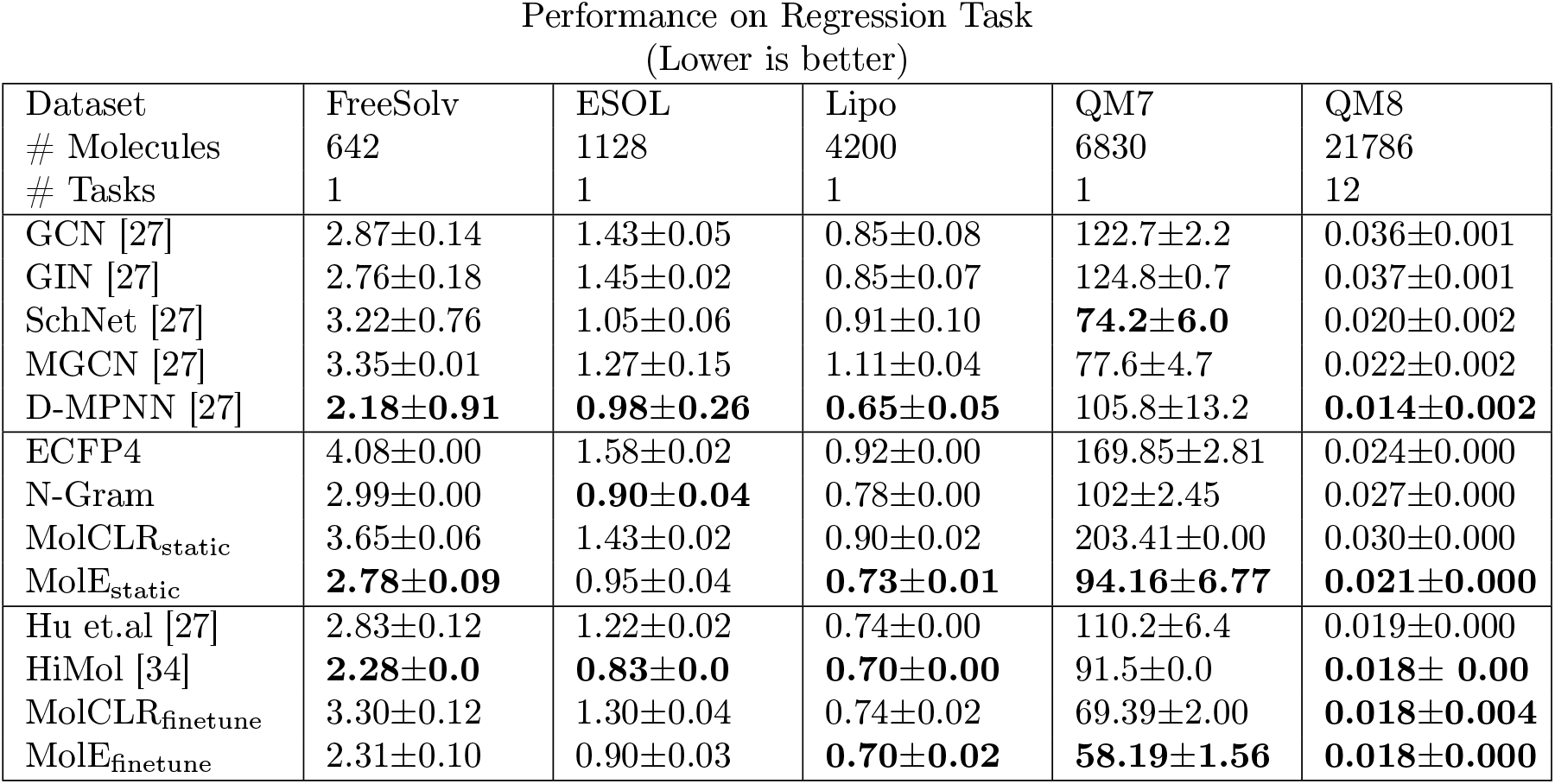
Average performance and standard deviation obtained on regression benchmark tasks. The first 5 models are supervised learning methods. The next 4 are the names of molecular features given as input to an XGBoost regressor. The final four methods are fine-tuned models. RMSE is shown for FreeSolv, ESOL, and Lipo. MAE is shown for QM7 and QM8. The best performance metric for each category is marked in **bold**.

**Extended Data Table 2:** Average ROC-AUC (%) and standard deviation obtained on classification benchmark tasks. The best performance metric is marked in **bold**

| Dataset<br># Molecules<br># Tasks | BBBP<br>2039<br>1 | Tox21<br>7831<br>12 | ClinTox<br>1478<br>2 | BACE<br>1513<br>1 | SIDER<br>1427<br>27 | HIV<br>41127<br>1 |
| --- | --- | --- | --- | --- | --- | --- |
| ECFP4 | 72.29 ± 1.36 | 68.67 ± 1.48 | 76.86 ± 2.66 | **84.53 ± 1.17** | 62.69 ± 2.38 | 76.86 ± 1.28 |
| N-Gram | 73.31 ± 0.36 | 73.96 ± 1.48 | **82.37 ± 3.5** | 80.58 ± 0.92 | 65.59 ± 2.13 | 75.52 ± 1.01 |
| MolCLR<sub>static</sub> | 69.01 ± 0.89 | 71.04 ± 1.93 | 76.32 ± 2.60 | 75.30 ± 1.96 | 64.12 ± 2.23 | 72.63 ± 1.09 |
| MolE<sub>static</sub> | **74.52 ± 1.24** | **75.98 ± 1.36** | 78.57 ± 7.10 | **84.60 ± 1.03** | **66.01 ± 2.49** | **78.34 ± 1.87** |

**Extended Data Table 3:** Average performance and standard deviation obtained on regression benchmark tasks. RMSE is shown for FreeSolv, ESOL, and Lipo. MAE is shown for QM7 and QM8. The best performance metric is marked in **bold**

| Dataset<br># Molecules<br># Tasks | FreeSolv<br>642<br>1 | ESOL<br>1128<br>1 | Lipo<br>4200<br>1 | QM7<br>6830<br>1 | QM8<br>21786<br>12 |
| --- | --- | --- | --- | --- | --- |
| ECFP4 | 4.23±0.00 | 1.66±0.00 | 0.95±0.001 | 147.16±0.73 | 0.024±0.000 |
| N-Gram | 2.89±0.02 | 1.13±0.00 | 0.82±0.001 | 95.35±0.60 | 0.029±0.000 |
| MolCLR<sub>static</sub> | 3.55±0.08 | 1.44±0.01 | 0.92±0.001 | 127.63±1.08 | 0.028±0.000 |
| MolE<sub>static</sub> | **2.80 ± 0.02** | **1.07 ± 0.01** | **0.79 ± 0.002** | **69.08 ± 0.11** | **0.022 ± 0.000** |

**Extended Data Figure 1:**
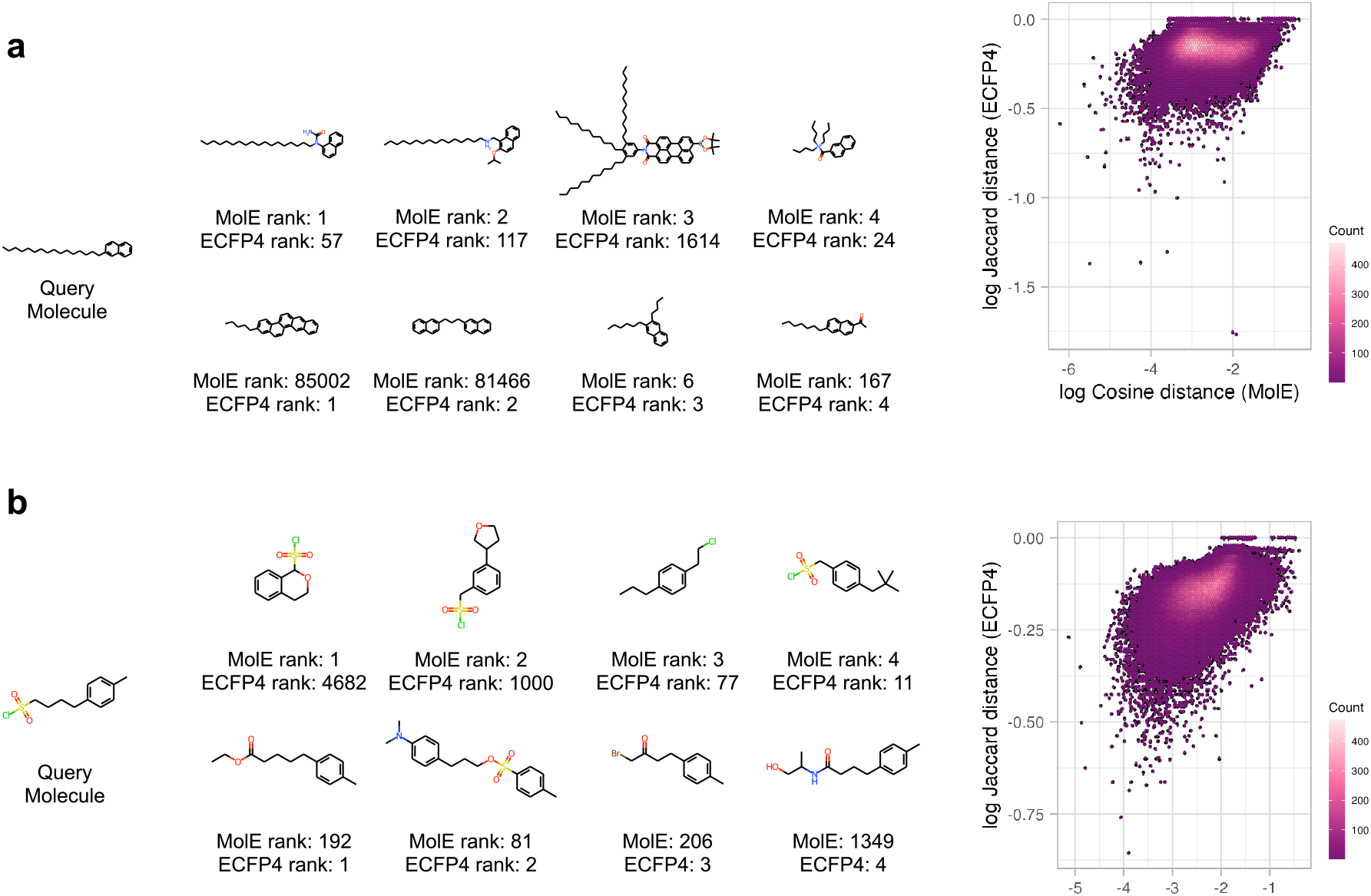
Comparing molecules most similar to a query as determined by ECFP4 and MolE. **a**. Highlighted similar structures and comparison of cosine and Jaccard distances estimated for PubChem ID: 12277389. **b**. Highlighted similar structures and comparison of cosine and Jaccard distances estimated for PubChem ID: 98701517.

**Extended Data Figure 2:**
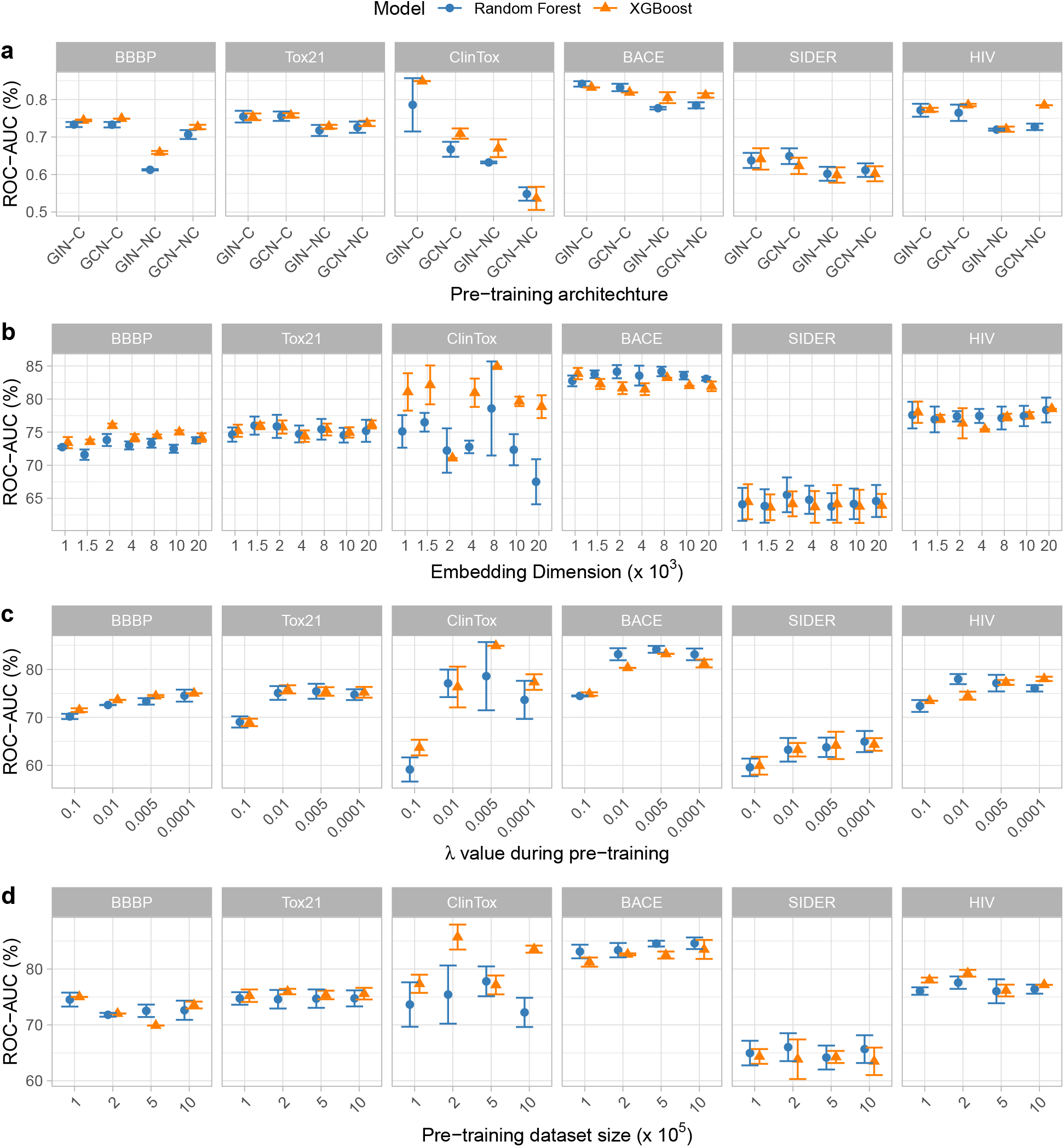
Pre-training ablation study. **a**. Performance when classifiers are trained on representations obtained with different pre-training architectures. **b**. Performance when classifiers are trained on representations obtained with different sizes of **z. c** Performance when classifiers are trained on representations obtained with different values of *λ*. **d** Performance when classifiers are trained on representations obtained after pre-training on different numbers of unlabeled molecular structures.

**Extended Data Figure 3:**
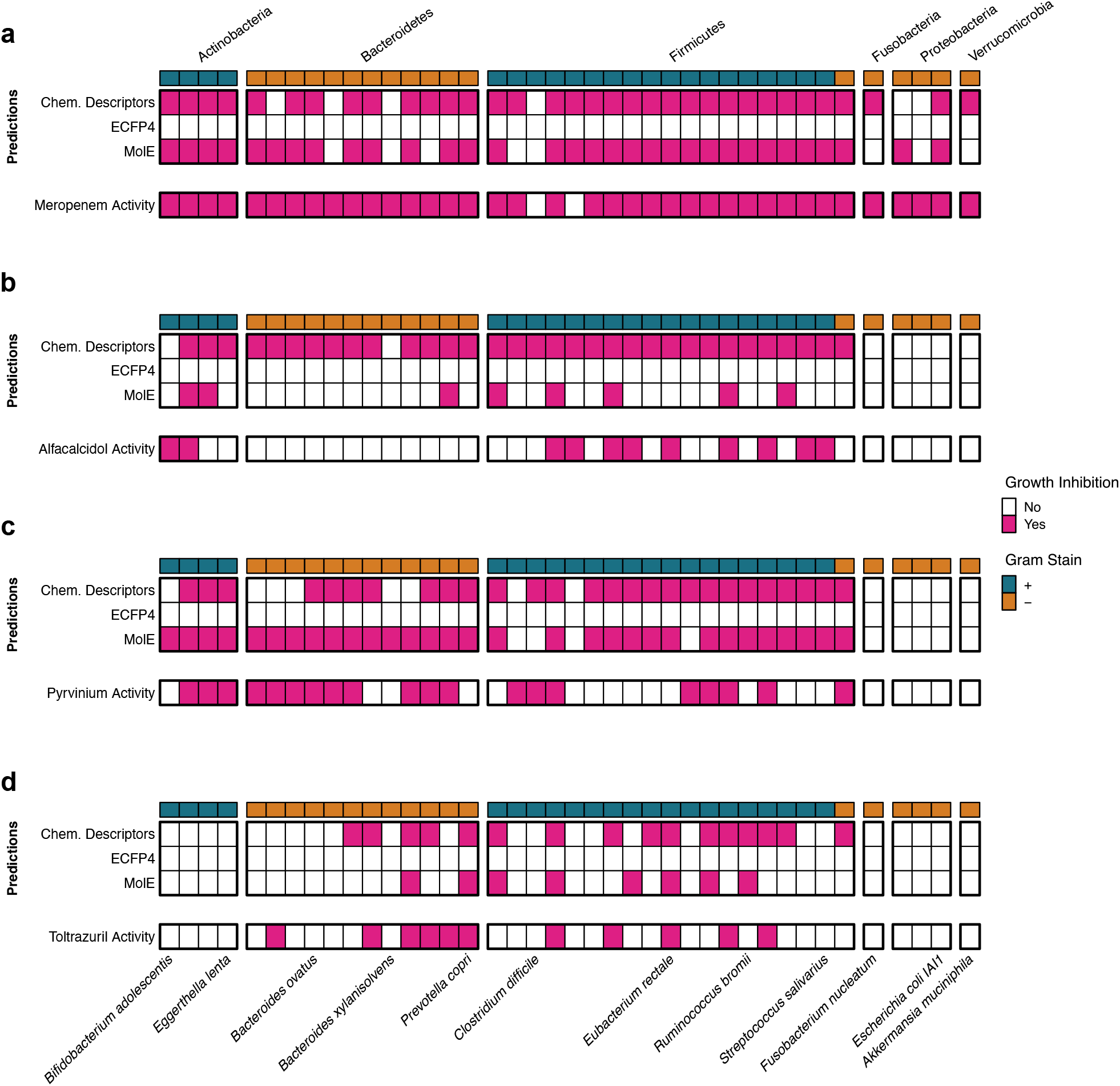
Examples of molecules in the test set that are missed by ECFP4 and recovered by MolE. Every panel shows the predictions made by models trained with chemical descriptors, ECFP4 and MolE are show in the first three rows, while the ground truth is shown in the bottom row. **a**. Meropenem, is a known antibiotic. **b**. Alfacalcidol is a vitamin D analog. **c**. Pyrvinium is an anti-parasitic medication. **d**. Toltrazuril is an antiparsitic drug.

**Extended Data Figure 4:**
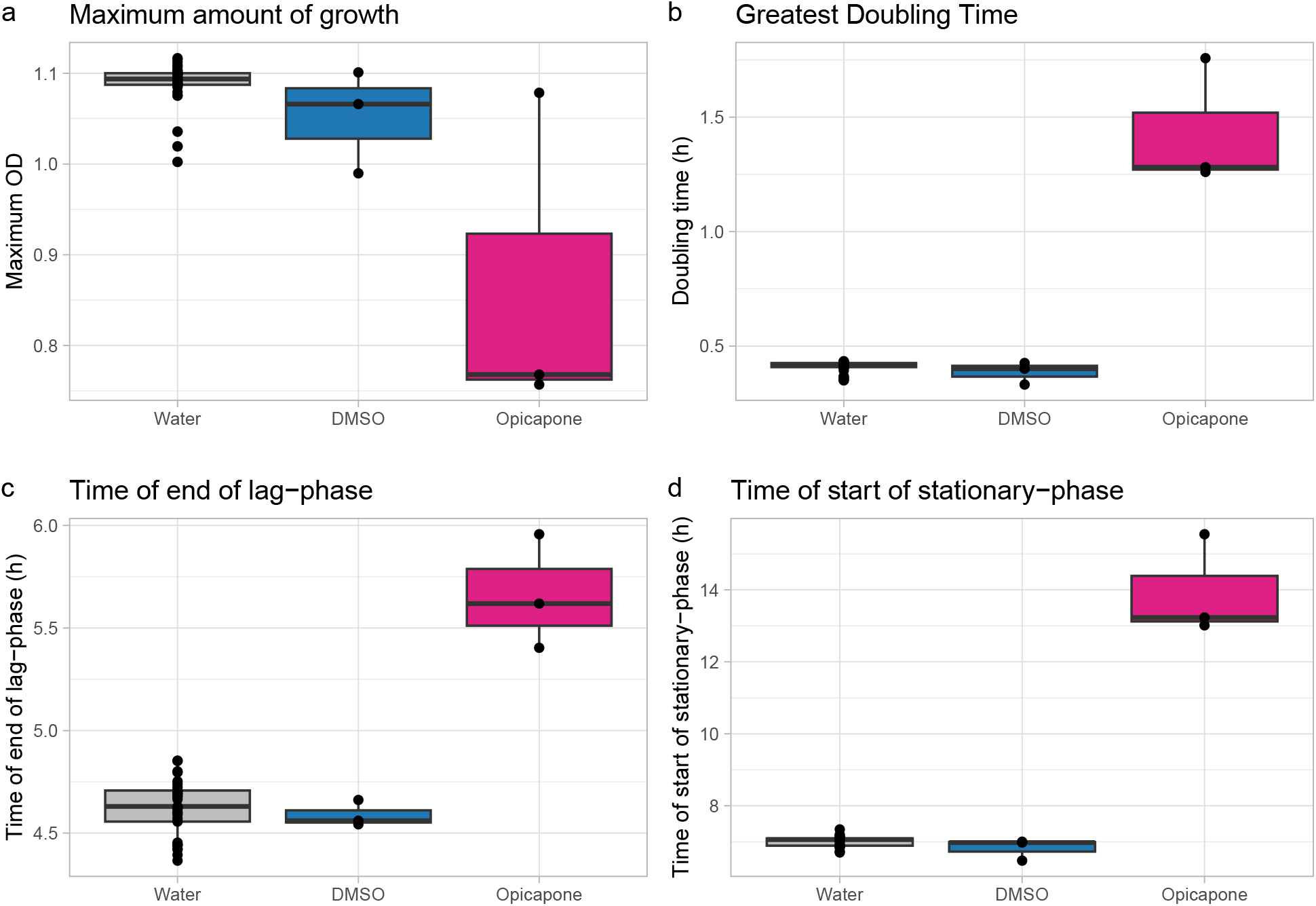
Modeled growth parameters for *S. aureus* under Opicapone (16 *µ*g/mL). **a**. Maximum modeled amount of growth (OD). **b**. Fastest doubling time (hours). **c**. Amount of time (hours) when lag-phase of growth ends. **d**. Amount of time (hours) when the stationary phase is reached.

**Extended Data Figure 5:**
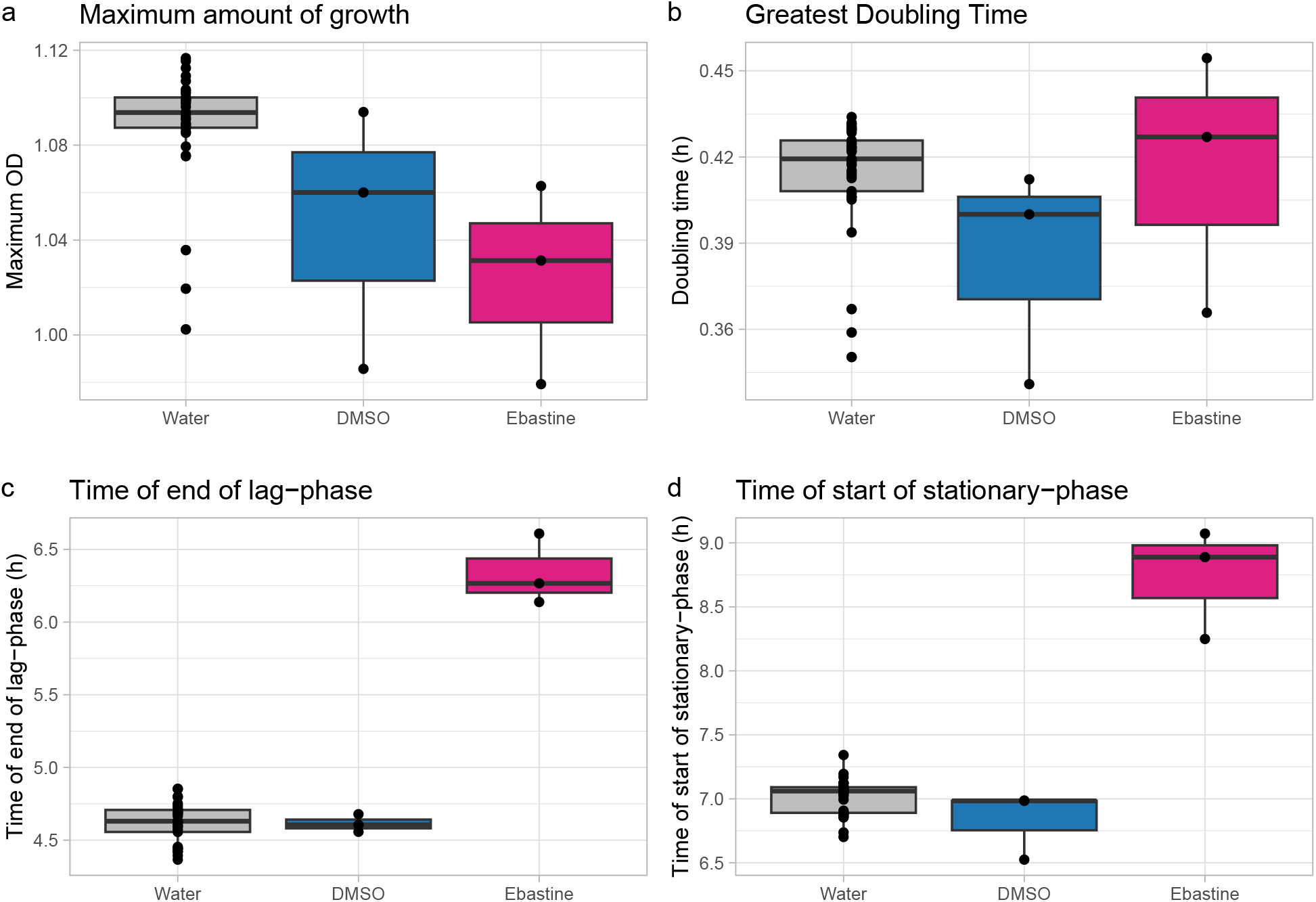
Modeled growth parameters for *S. aureus* under Ebastine (16 *µ*g/mL). **a**. Maximum modeled amount of growth (OD). **b**. Fastest doubling time (hours). **c**. Amount of time (hours) when lag-phase of growth ends. **d**. Amount of time (hours) when the stationary phase is reached.

**Extended Data Figure 6:**
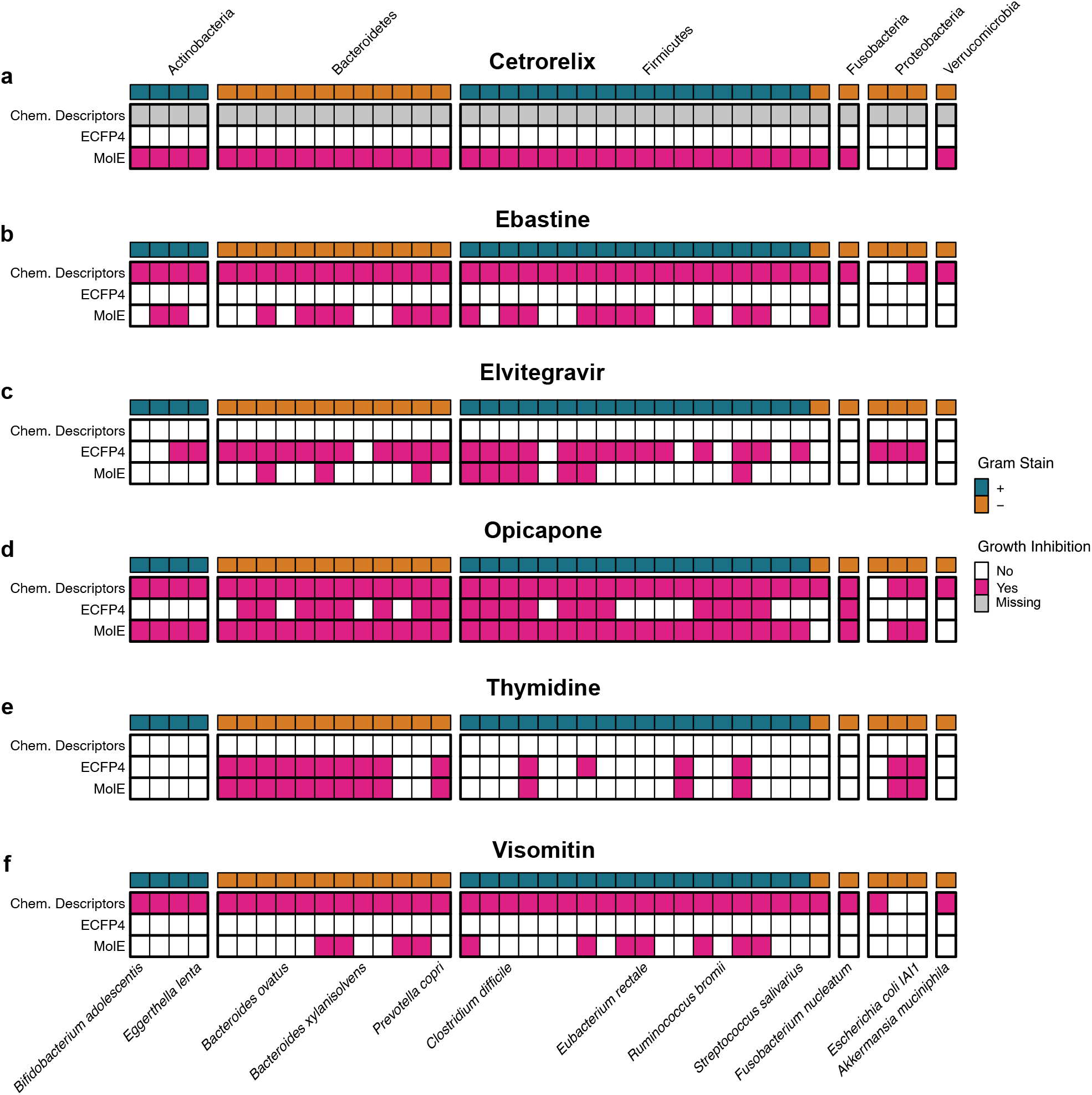
Predicted antimicrobial activity of compounds selected for experimental validation. a. Cetrorelix. No chemical descriptors were able to be estimated for this compound, therefore no predictions were made. **b**. Ebastine. No antimicrobial activity was predicted when using ECFP4 features. **c**. Elvitegravir. No antimicrobial activity was predicted when using chemical descriptors as features. **d**. Visomitin. No antimicrobial activity was predicted when using ECFP4 features.

