## Supplementary material for "Pre-trained molecular representations enable antimicrobial discovery": Supplmentary Information

Random Forest Parameters

| Name | Possible Values |
|---|---|
| criterion | "gini", "entropy", "squared_error"* |
| max_features | "sqrt", "log2", None |
| n_estimators | 100, 300, 500, 700, 1000 |
| class_weight+ | "balanced", None |

Supplementary Table 1: Random Forest parameters considered during a random search. Some parameters are only considered during *regression or +classification tasks

XGBoost Parameters

| Name | Possible Values |
|---|---|
| max_depth | 5, 10, 50, 100 |
| eta | 0.3, 0.1, 0.05, 1 |
| n_estimators | 30, 100, 300, 500, 1000 |
| subsample | 0.3, 0.5, 0.8, 1.0 |

Supplementary Table 2: XGBoost parameters considered during a random search.

Finetuning parameters

| Name | Possible Values |
|---|---|
| Batch size | 32, 100, 512, 800 |
| Learning rate for MLP head | 5 x $10^{-4}$, $10^{-3}$ |
| N layers in MLP head | 1, 2 |
| Activation function MLP | "softplus", "relu" |
| Learning rate for GNN | 5 x $10^{-5}$, $10^{-4}$, 2 x $10^{-4}$, 5 x $10^{-4}$ |
| Dropout rate | 0, 0.1, 0.3, 0.5 |

Supplementary Table 3: Finetuning parameters considered during a random search.

Compounds selected for experimental validation

| Name | Summary | Structure |
|---|---|---|
| Cetrorelix | A synthetic peptide antagonist of gonadotropin-releasing hormone used to prevent luteinizing hormone surges in women undergoing assisted reproduction therapy | 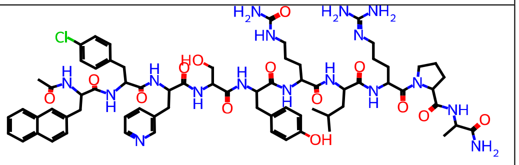 |
| Ebastine | A second generation H1-receptor antagonist useful in the treatment of allergic rhinitis and urticaria. | 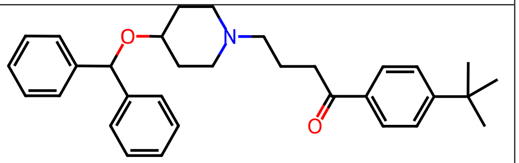 |
| Elvitegravir | An antiretroviral agent used for the treatment of HIV-1 infection. | 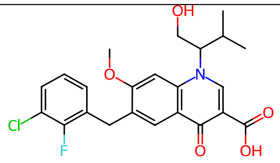 |
| Opicapone | A third-generation catechol-O-methyltransferase inhibitor used as an adjunct treatment for Parkinson's Disease. | 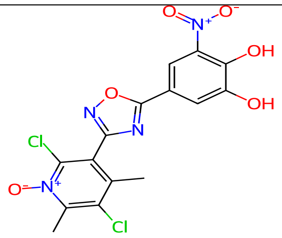 |
| Thymidine | A specific precursor of deoxyribonucleic acid is used as a cell-synchronizing agent. | 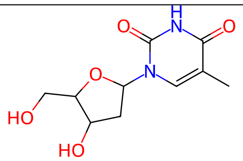 |
| Visomitin | A mitochondrial-targeted antioxidant with a high mitochondrion membrane penetrating ability and potent antioxidant capability. | 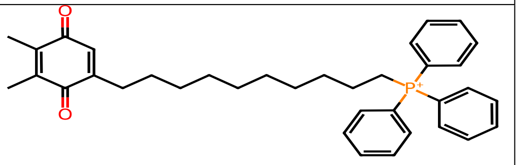 |

Supplementary Table 4: Compounds selected for experimental validation. Summaries were gathered from DrugBank [1] and MedChemExpress

Bacterial strains used for experimental validation

| Strain | Source | Notes |
|---|---|---|
| *Escherichia coli* IAI1 | Nassos Typas | Commensal strain |
| *Escherichia coli* CFT0731 | DSMZ | UPEC model strain (`https://www.dsmz.de/collection/catalogue/details/culture/DSM-103538`) |
| *Pseudomonas aeruginosa* PA14 | Nassos Typas | Domesticated clinical isolate |
| *Staphylococcus aureus* subsp. str. Newman | Nassos Typas | NCTC 8178; ATCC: 13420; `http://www.lgcstandards-atcc.org/Products/All/25904.aspx` |
| *Klebsiella pneumoniae* MKP103 | Nassos Typas | Lab strain |

Supplementary Table 5: . Bacterial strains screened during experimental validation
